# Dynamic rewiring of biological activity across genotype and lineage revealed by context-dependent functional interactions

**DOI:** 10.1101/2021.06.25.450004

**Authors:** Eiru Kim, Lance C. Novak, Veronica Gheorghe, Christopher A. Bristow, Traver Hart

## Abstract

Coessentiality networks derived from CRISPR screens in cell lines provide a powerful framework for identifying functional modules in the cell and for inferring the role of uncharacterized genes. However, these networks integrate signal across all underlying data, and can mask strong interactions that occur in only a subset of the cell lines analyzed. Here we decipher dynamic functional interactions by identifying significant cellular contexts, primarily by oncogenic mutation, lineage, and tumor type, and discovering coessentiality relationships that depend on these contexts. We recapitulate well-known gene-context interactions such as oncogene-mutation, paralog buffering, and tissue-specific essential genes, show how mutation rewires known signal transduction pathways, including RAS/RAF and IGF1R-PIK3CA, and illustrate the implications for drug targeting. We further demonstrate how context-dependent functional interactions can elucidate lineage-specific gene function, as illustrated by the maturation of proreceptors *IGF1R* and *MET* by proteases *FURIN* and *CPD*. This approach advances our understanding of context-dependent interactions and how they can be gleaned from these data. We provide an online resource to explore these context-dependent interactions at diffnet.hart-lab.org.

## Introduction

Development of genome-wide CRISPR screening half a decade ago facilitated robust determination of proliferation dependent genes for each cancer type (Hart et al., 2015; Shalem et al., 2014; Wang et al., 2015). Current efforts for identifying cancer vulnerability in public are done for hundreds of cancer cell lines (Aguirre et al., 2016; Behan et al., 2019; Meyers et al., 2017) across dozens of tissue types. Previously, we and others demonstrated that functionally coherent modules of genes can be extracted from CRIPSR screens by measuring the similarity of gene knockout fitness profiles across all gene pairs, generating a functional interaction network based on coessentiality (Boyle et al., 2018; Kim et al., 2019; Pan et al.; Rauscher et al., 2018; Wainberg et al., 2021; Wang et al., 2017). Many such modules found in the network encode well annotated biological processes, enabling robust functional prediction for uncharacterized genes (Aregger et al., 2020; Noordermeer et al., 2018; Zimmermann et al., 2018). Some modules, however, are associated with specific tissues or genomic perturbations, such as the *ESR1*-*FOXA1* pathway in ER+ breast cancer cell lines and the *BRAF*^*V600E*^ module in melanoma cells (Amici et al., 2020; Kim et al., 2019; Sharma et al., 2020). In this latter cluster, gene mutation is so conflated with tissue specificity that b-RAF is tightly linked with melanocyte-specific transcription factors *MITF* and *SOX9* but does not show significant correlation with its known protein interaction partner c-RAF (encoded by the *RAF1* gene).

We hypothesized that mutations in specific genes, and more broadly the genetic and epigenetic state of cells, could rewire these coessentiality networks, and that disentangling this rewiring could elucidate the biological and therapeutic implications of genetic lesions and tissue-specific diseases. Context-specific interaction rewiring or differential network analysis is a useful tool for understanding how fixed genomes can emerge into heterogeneous cellular and tissue morphologies and phenotypes (Ding et al., 2018; Ideker and Krogan, 2012; Kim et al., 2012). Previously, studies of comparison of coexpression between two contexts selected from a pool were introduced to discover differentially regulated interactions by specific context (Cho et al., 2009; Hsu et al., 2015; Lui et al., 2015). In the pre-CRISPR era, integrative functional interaction networks were exploited to prioritize disease genes (Kim et al., 2008; Lee et al., 2004; Singh-Blom et al., 2013), and included efforts to identify tissue-specific networks (Greene et al., 2015; Guan et al., 2012), but these approaches lack the power of inferred genetic interaction profiles that coessentiality networks derived from screens in human cells provide. Recent examinations of coessentiality networks such as CEN-tools (Sharma et al., 2020) and fireworks (Amici et al., 2020) offer the ability to browse how correlates of a query gene vary by background, but offer little ability to interpret the results. Moreover, the ability to go from lesion of interest to lesion-associated network rewiring is both limited and difficult to use.

In this study, we developed a framework of identifying dynamic relationships in cancer dependency data caused by functional contexts such as variations in tissue of origin and/or genomic lesions. We first categorized genomic perturbations using molecular profiles from the Cancer Cell Line Encyclopedia. Then, we investigated which genomic features were associated with synthetic gene essentiality by applying a logistic regression model, and used these features to search for associated network rewiring. Rewired networks carry information on drug efficacy, the biological impact of specific lesions, and the tissue-specific activity of genes.

## Results

### Associating cellular context with emergent gene essentiality

Understanding the causal, or at least associative, basis of variation in gene essentiality is important in matching the right anti-cancer drugs with responsive tumors (Rancati et al., 2018). For example, KRAS is essential when it undergoes oncogenic mutation (Janes et al., 2018; O’Leary, 2021; Waters and Der, 2018), and SWI/SNF complex member ARID1B is essential if its paralog ARID1A is deleted (Helming et al., 2014). To globally decipher these essential gene/context relationships, we first systematically categorized genomic lesions into gain of function/hotspot (GOF) or loss of function (LOF) using mutation data from 808 cell lines in the Cancer Cell Line Encyclopedia (Barretina et al., 2012) that have matching CRISPR screen data from the Cancer Dependency Map (Behan et al., 2019; Meyers et al., 2017; Tsherniak et al., 2017). We added cell line metadata including lineage/tumor type, as well as a feature describing EMT state based on the ratio of *CDH1* to *VIM* expression (see Methods). After de-duplication and filtering, we retained 2,918 binary features as predictor variables.

We used a machine learning approach to estimate the effect of genomic perturbations on variably essential genes (**Figure 1A**). We chose logistic regression with an elastic net penalty because it handles binary predictors, because gene essentiality is readily binarized for use as a response variable, and because the L1 component of the elastic net forces most coefficients to zero, resulting in a model that is easily interpreted. For each gene, we generated a binary essentiality vector across 808 good quality cell lines in DepMap 20Q4 data processed with our BAGEL2 pipeline (Kim and Hart, 2020) for use as a response variable. We used logistic regression on the same feature table to predict the essentiality profile of each gene independently. This approach is similar to that used by Lord et al (Lord et al., 2020), but our logistic regression is a binary classifier of gene essentiality rather than their linear regression on continuous fitness effects.

**Figure 1.**
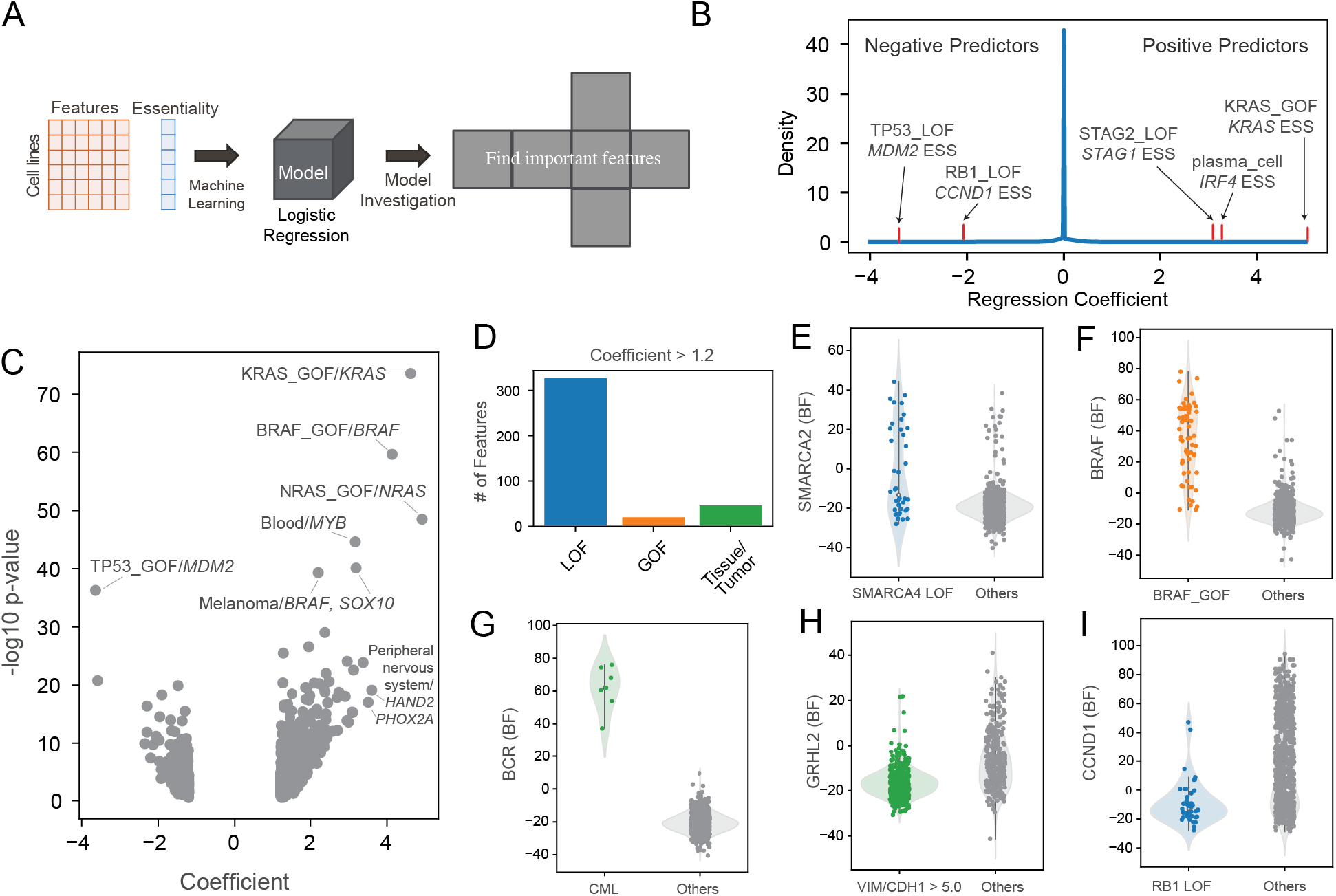
Underlying drivers of gene essentiality. **A)** a schematic process of finding important contexts for a gene. We input binary feature vectors across all cell lines and a vector of essentiality of a gene and trained a logistic regression model. Then, we investigated coefficients of features to determine the features important for predicting gene essentiality. **B)** Distribution of regression coefficients collected from prediction models of all genes. Positive coefficients reflect gene essentiality in the presence of the feature, and a negative coefficients reflect increased gene essentiality in the absence of the feature. **C)** Volcano plot demonstrated a general correlation between P-values of Fisher exact test (Y axis) and regression coefficients (X axis). **D)** Category of 393 features with |regression coefficient| > 1.2. **E-I)** Examples of predictions from **E)** paralog interaction, **F)** oncogene addiction, **G)** tissue specific genes, **H)** EMT expression signature, and **I)** negative association. BF, Bayes Factor from BAGEL2 analysis of DepMap data (positive=essential).

After calculating all regression models, we investigated which genomic features significantly contributed to predicting essentiality (**Figure 1B,C**). We collected coefficients of genomic features across all prediction models of genes, which loosely represent the magnitude of predictive power. A positive coefficient of a genomic feature indicates that the genomic perturbation is associated with the gene dependency (e.g. KRAS_GOF/*KRAS* essentiality), and a negative coefficient indicates that gene essentiality is associated with the absence of the lesion (e.g. TP53_LOF/*MDM2* essentiality). Coefficients showed a similar trend with P-values derived by the Fisher exact test between a vector of genomic lesions and a vector of gene dependency across cell lines (Figure 1C). By considering strong coefficients (see Methods and **Supplementary Figure 1**), we identify 393 genomic features strongly predictive of variation in gene essentiality, including 327 loss of function mutations, 20 gain of function mutations, and 46 features derived from cell metadata (**Figure 1D**). We recapitulated positive associations in paralog buffering (e.g. SMARCA4_LOF/*SMARCA2*^*ess*^, **Figure 1E**), oncogenic mutation-induced essentiality (e.g. BRAF_GOF/*BRAF*^*ess*^, **Figure 1F**),, and tumor-specific dependencies (e.g. CML/*BCR*^*ess*^, **Figure 1G**), as well as negative associations such as epithelial transcription factor *GRHL2* essentiality in cells with low *CDH1/VIM* expression ratio (**Figure 1H**) and increased essentiality of cyclin D1 (*CCND1*) in the absence of downstream *RB1* loss of function (**Figure 1I**). These results are highly concordant with previous predictive models (**Supplementary Figure 1**) (Behan et al., 2019; Lord et al., 2020) and known biology, and provide a meaningful set of features for exploring context-dependent functional interaction rewiring.

### Measuring context-dependent network rewiring

With the identification of contexts associated with variation in gene essentiality, we expanded our view of dynamics to functional interaction rewiring caused by these genomic and epigenomic features. In general, genes with correlated fitness profiles across a diverse panel of cell lines tend to operate in the same biological processes, such as enzymatic or signal transduction pathways, often as subunits in the same protein complex. While this correlation provides a powerful framework for inferring gene function and the modular structure of the cell (Boyle et al., 2018; Kim et al., 2019; Pan et al.; Rauscher et al., 2018; Wainberg et al., 2021; Wang et al., 2017), it relies on covariation in the underlying gene essentiality vectors but offers no insight into the source of this variation. We hypothesized that, by systematically removing rationally-selected subsets of cell lines and measuring the effect on pairwise correlation, we could identify the drivers of this covariation and, in turn, infer the causal basis for the emergent essentiality of biological processes.

Although differential co-essentiality offers a seemingly straightforward way to approach context rewiring of functional interaction networks, the method is beset by complications. One is that, for many genes, physiologically relevant knockout fitness defects are typically only seen in a few cell lines. A related issue is that there are frequently too few cell lines associated with a given context for correlation to be an accurate predictor of functional interaction within the context; e.g. a CML-only coessentiality network has limited meaning if there are only seven CML cell lines. To overcome these difficulties, we designed a strategy comprising a leave-one-out test and bootstrapped network comparison analysis to investigate snapshots of interaction rewiring in essentiality data associated with our features of interest. In the leave-one-out test, interaction rewiring was measured as a differential Pearson correlation coefficient (dPCC) of a context by taking the PCC from all cell lines (PCC_all_) and subtracting the PCC using all cell lines except those carrying the feature of interest (PCC_∼context_) (**Figure 2A**). We classified a positive dPCC as a gain of interaction (GOI) because the PCC depends on the presence of cells with the feature of interest (**Figure 2A**, top). Conversely, we describe a negative dPCC as a loss of interaction (LOI) because the background PCC is improved by removing cells with the feature of interest (**Figure 2A**, bottom).

**Figure 2.**
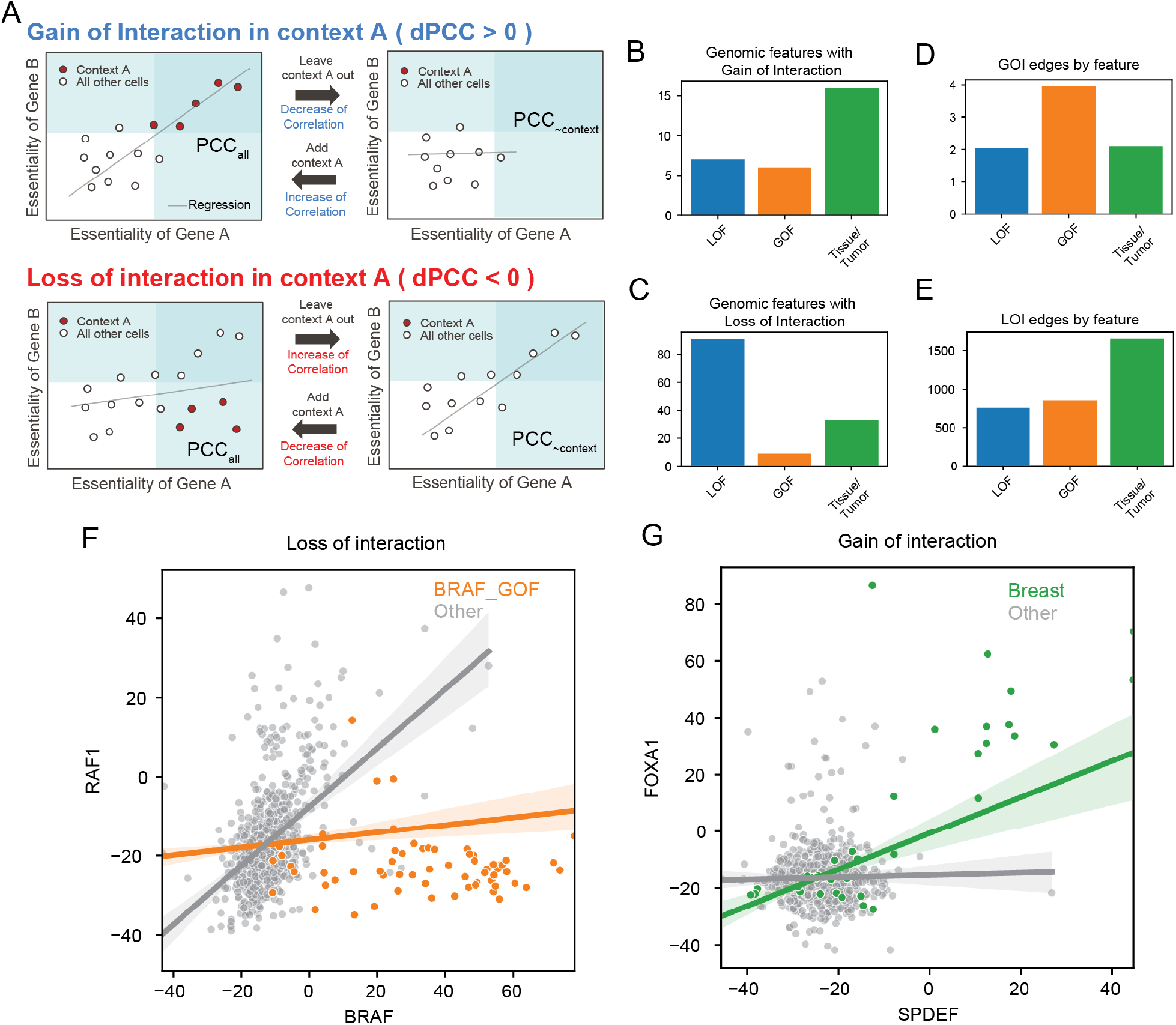
A framework to identify dynamics in coessentiality network. **A)** Classes of context-dependent interaction. In all cases, a differential Pearson correlation coefficient (dPCC) for each gene pair is calculated as the difference between the global network (PCC_all_) and the PCC from all cells except those harboring the feature of interest (PCC_∼context_). **B,C)** The number of features associated with gain or loss of interaction. **D,E)** the number of gained or lost interaction, by feature type. **F-G)** Examples of context-dependent loss and gain of interaction. Axes are Bayes Factors of the genes.

We measured dPCC for the 393 features identified in logistic regression models as significant predictors of gene essentiality, and measured significance by bootstrap resampling to estimate a null distribution of dPCC (See Methods and **Supplementary Figure 2**). We identified 30 features associated with gain of interaction (**Figure 2B**) and 133 features, including 91 LOF mutants, associated with loss of functional interaction (**Figure 2C**) in the cell lines studied. These features are associated with 9,227 gain of interaction and 3,261 loss of interaction events (**Figure 2D-E**). Notably, of the gain of interaction events, 97% (n=8,958) are associated with TP53 gain of function/hotspot mutations, leaving only 269 interactions associated with the remaining 29 features.

Two case studies illustrate the effect of context-dependent rewiring of functional interaction networks. RAS and RAF kinases are the core members of RTK signaling pathways for proliferation and differentiation (Asati et al., 2016; Lavoie and Therrien, 2015; Santarpia et al., 2012; Terrell and Morrison, 2019; Terrell et al., 2019). Within the canonical MAP kinase signal transduction pathway, b-RAF (*BRAF*) and c-RAF (*RAF1*) form a heterodimer to transmit phosphorylation signals from upstream RAS genes to downstream MEK/ERK targets. In *BRAF*-driven cancers, oncogenic mutation constitutively activates *BRAF* and obviates the need for c-RAF binding. In the global coessentiality network, *BRAF*^*V600E*^ and *BRAF*^*wt*^ cell lines are mixed, diluting the ability to discover the wildtype *BRAF*-*RAF1* relationship (background PCC=0.124). Removing *BRAF*^*V600E*^ cell lines boosts this correlation to 0.478 (dPCC = −0.363; P<0.001, permutation test); thus, BRAF_GOF mutation is causal for a loss of interaction between *BRAF* and *RAF1* (**Figure 2F**). Similarly, transcription factor *FOXA1* is essential in a subset of lung and breast cancers but shows a breast-specific interaction with ETS family transcription factor *SPDEF* (**Figure 2G**).

### Reconstruction of Directionality in Signaling Pathways

The dynamic rewiring of functional interactions resulting from cancer-associated mutation has predictable consequences. As previously observed, mutation of the *TP53* tumor suppressor removes the cell’s reliance on *MDM2* and related genes to suppress the proapoptotic activity of wildtype p53 protein. Similarly, constitutive activation of signaling proteins by oncogenic mutation activates downstream signaling partners while removing the need for upstream activation signals. Here we describe two such pathways.

#### RAS/RAF signaling pathway

Mutations in the RAS pathway are major drivers of numerous cancers. RAS family members *KRAS* and *NRAS* are frequently mutated in colorectal and pancreas adenocarcinoma, and multiple myeloma and melanoma, respectively. *BRAF* is a major driver gene of melanoma and thyroid adenocarcinoma (Gonzalez-Perez et al., 2013; Martínez-Jiménez et al., 2020). Oncogenic GOF lesions in these genes caused significant intra- and inter-pathway interaction rewiring (**Figure 3A**). *KRAS* and *NRAS* are usually non-essential in CRISPR data; however, mutation or amplification of these genes induces hyperactivation and strong mutually exclusive essentiality. We found that Interaction rewiring of GOF mutations of *KRAS, NRAS*, and *BRAF* induced gain of interaction with downstream genes and disconnected the network from upstream genes through loss of interaction (Figure 3A). For example, *KRAS* mutation amplifies the link with downstream signaling partner *RAF1* (PCC_all_ = 0.390, PCC_∼KRAS_GOF_ = 0.239, dPCC = 0.151, P<0.001), severs the link between *RAF1* and *KRAS* paralog *NRAS* (PCC_all_ = 0.305, PCC_∼KRAS_GOF_ = 0.472, dPCC = −0.167, P<0.001), and disconnects MAP kinase signaling from the upstream *EGFR* receptor tyrosine kinase signal transduction module (*KRAS-SOS1* PCC_all_ = −0.012, PCC_∼KRAS_GOF_ = 0.184, dPCC = −0.197, P<0.001). Similar behavior is seen in *NRAS* and *BRAF* GOF cell lines (**Figure 3B-E**). Additional rewiring by *BRAF* mutation abrogates interaction with dimerization partner *RAF1*, consistent with the known biology of *BRAF* mutation. We found that *BRAF* mutation induced loss of interaction between *BRAF* and *RAF1* while *BRAF* and *RAF1* are well correlated in wildtype *BRAF* cell lines (**Figure 3A,C**).

**Figure 3.**
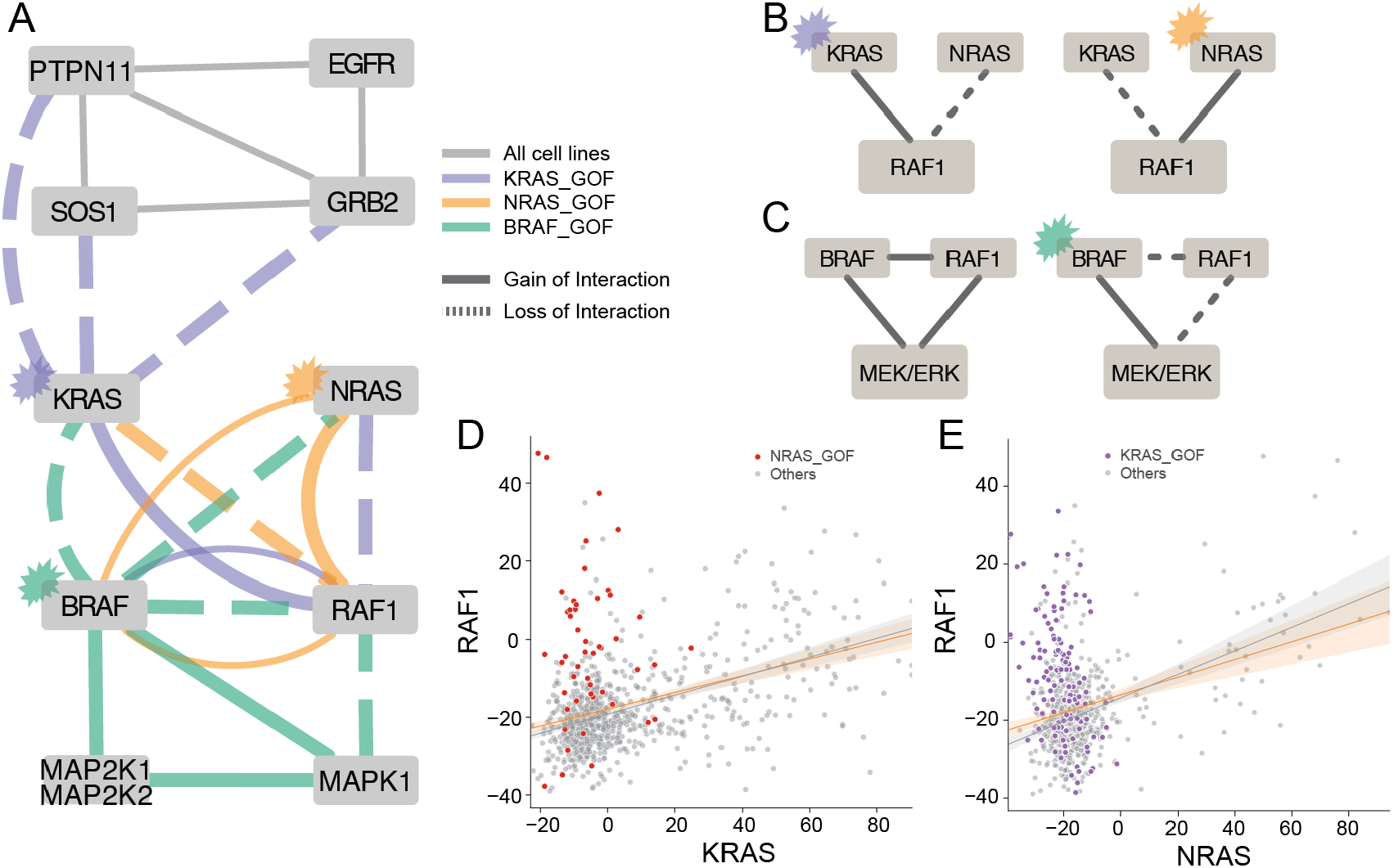
Interaction rewiring in RAS/RAF pathway. **A)** a comprehensive network diagram of differential interactions caused by mutations on KRAS, NRAS, and BRAF. **B)** KRAS mutant caused loss of NRAS-RAF1 interaction and NRAS mutant caused loss of KRAS-RAF interaction. **C)** RAF1 is required for regulating MEK/ERK pathway when BRAF is wildtype, while BRAF mutant doesn’t require RAF1 and led to loss of interaction of RAF1 to BRAF and the downstream pathway. **D,E)** Scatter plots of interactions between RAF1 and two RAS proteins.

#### IGF1R/PIK3CA signaling pathway

The *PIK3CA* gene encodes a protein kinase that mediates signaling from insulin like growth factor receptor *IGF1R* to the *AKT* pathway (Zha and Lackner, 2010) (**Figure 4A**), is frequently mutated in a number of cancers, and is itself the target of several chemotherapeutic agents. When mutated, *PIK3CA* is hyperactivated and induces downstream pathways without reliance on *IGF1R* signals (Gkeka et al., 2014). In our differential network analysis, *PIK3CA* gain of function mutation causes not only loss of interaction with *IGF1R* receptor (**Figure 4B**) but also the insulin receptor substrate gene *IRS2*, which encodes a protein involved in *IGF1R* signal transduction (**Figure 4C**). In addition, PIK3CA_GOF causes loss of interaction with *FURIN* (**Figure 4D**), a protease required for maturation of the *IGF1R* receptor (*FURIN*-*IGF1R* PCC_all_= 0.540, n=808 cell lines) and uncharacterized gene *KBTBD2* (**Figure 4E**). The high *KBTBD2*-*IRS2* background correlation (PCC_all_ = 0.483), and the loss of interaction with *PIK3CA* upon oncogenic mutation (PCC_all_ = 0.293, PCC_∼PIK3CA_GOF_ = 0.400, dPCC = −0.107, P<0.001), support a functional interaction between *KBTBD2* and *IRS2*, possibly involving protein maturation or signaling.

**Figure 4.**
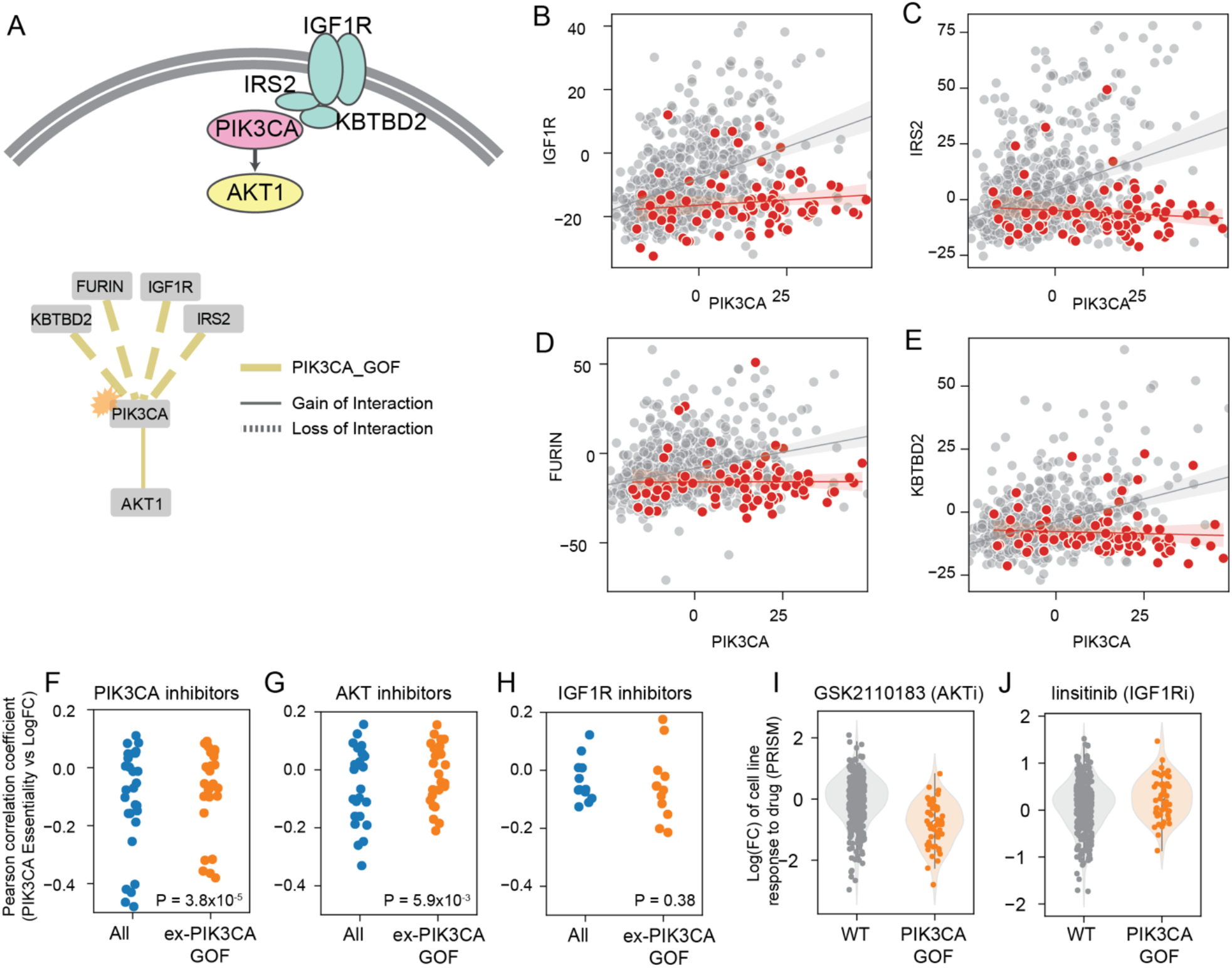
Dynamics of network rewiring on IGF1R-PIK3CA pathway. A) A visualization of IGF1R-PIK3CA pathways (upper panel) and a differential network of pathway genes caused by PIK3CA GOF mutation. Upstream pathway genes were disrupted by mutation while correlation with downstream pathway gene AKT1 was boosted. **B-E**) Scatter plots of interactions of PIK3CA upstream genes **B)** IGF1R and **C)** IRS2, plus **D)** IGF1R maturation factor FURIN, and **E)** KBTBD2. **F-H**) Comparisons of effectiveness of **F)** PIK3CA inhibitors, **G)** AKT inhibitors, and **H)** IGF1R inhibitors in all cells (blue) and cells excluding PIK3CA GOF mutations (orange), from PRISM. **I-J**) AKT inhibitor is more effective in PIK3CA GOF, while IGF1R inhibitor is more effective in PIK3CA wt.

We further investigated the impact of interaction rewiring on drug efficacy. We compared the effect of drugs targeting *PIK3CA, AKT1*, and *IGF1R*, between wild-type and mutant *PIK3CA* cell lines in PRISM (Yu et al., 2016) 19Q3 data (**Figure 4F-H**), where strong drug efficacy is shown as negative log fold change. Cell lines harboring a *PIK3CA* oncogenic mutation were significantly more sensitive to AKT inhibitor GSK2110183, (**Figure 4I**). Conversely, *IGF1R* inhibitor linisitib is effective only in a subset of cell lines with wildtype *PIK3CA* alleles (**Figure 4J**). Together, these findings are consistent with the mutational rewiring of the functional interaction network and demonstrate that context-dependent networks can inform drug sensitivity.

### Tissue-specific functional interaction

While *IGF1R* is associated with its canonical downstream signaling partner *PIK3CA*, its strongest association in the global coessentiality network is with *FURIN* (PCC_all_ = 0.540), reflecting the role of *FURIN* in *IGF1R* receptor maturation. *FURIN* is also strongly associated with Carboxypeptidase D (*CPD*), recently reported to be required for pro-*IGF1R* maturation in lung adenocarcinoma (Alarcón et al., 1994; Komada et al., 1993) (**Figure 5A**). Interestingly, the link between *CPD* and *IGF1R* is largely abrogated in glioma cells (**Figure 5B**), while the *CPD*-*FURIN* relationship is maintained (**Figure 5C**). Together these observations suggest that the *CPD*-*FURIN* activity is required for maturation of a different protein in glioma cells. Notably, receptor tyrosine kinase *MET* shows a strong gain of interaction with *CPD* in glioma (**Figure 5D**). *MET* polypeptide, like *IGF1R*, is proteolytically processed into a mature receptor through cleavage into alpha and beta subunits and linkage via disulfide bonds. Our context-dependent coessentiality network therefore provides corroborating evidence in support of *FURIN*/*CPD* processing of *MET* as suggested by Han *et al*. (Han et al., 2020).

**Figure 5.**
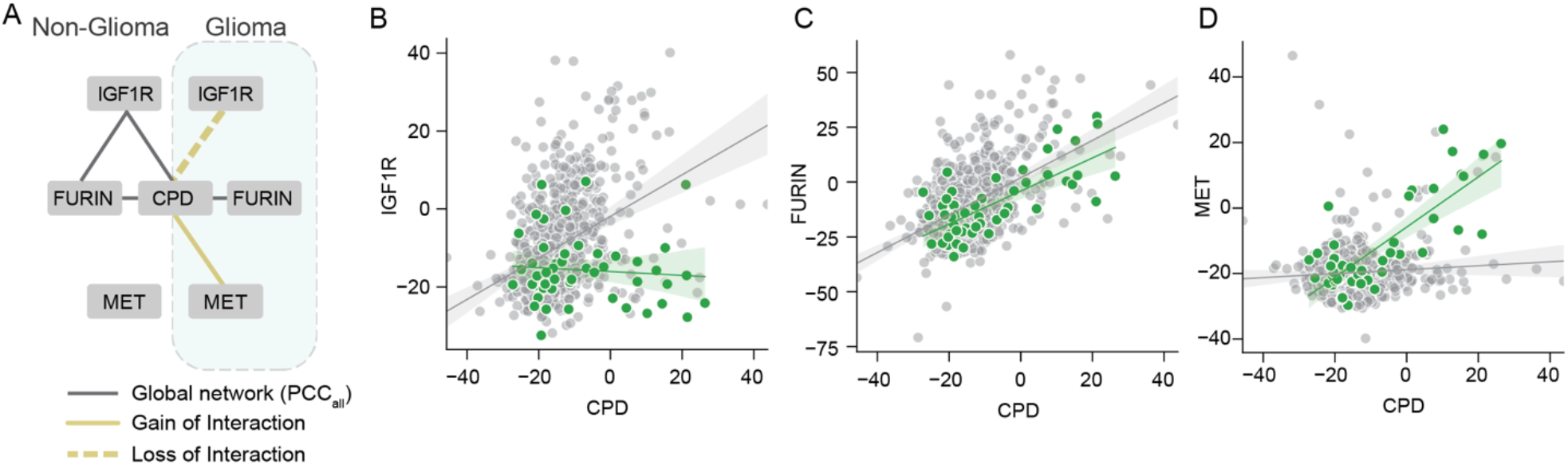
Tissue-specific network rewiring. **A)** A network diagram of interaction rewiring of CPD in glioma cell lines. **B)** IGF1R-CPD interaction was generally correlated in non-glioma cell, but loss of interaction was detected in glioma cell lines. **C)** CPD-FURIN interaction was correlated regardless of tissue type. **D)** Instead of IGF1R, MET is correlated with CPD in glioma cell lines.

## Discussion

In this study, we present a framework for understanding the differential wiring of cells across genotypes and lineages. Using genetic mutation information and cell line metadata, we investigated contexts that cause emergent essentiality by analyzing coefficients obtained from a logistic regression model trained using CRISPR screen data. Using this approach, we identified numerous context-gene interactions, including known interactions such as paralog buffering, oncogenes, and tissue specific essential genes.

We then examined the rewiring of functional interaction networks in the presence of these contexts. We developed a strategy which compares the strength of interaction between two genes, as measured by the Pearson correlation coefficient of their normalized fitness vectors across all samples, to the correlation derived from samples that exclude a specified context, or a “leave-one-out” test. We measured the significance of these changes by bootstrapping a null distribution for every gene pair in every context. Rewiring detected by our approach showed concordance with biological knowledge and discovered new putative context-dependent functional interactions, demonstrating its potential for functional genomics and cancer targeting. We provide a web-based interface for exploring the network edges rewired by mutation and/or lineage at diffnet.hart-lab.org.

The approach derived herein attempts to address one of the key questions arising from the use of coessentiality, or indeed any gene similarity approach, to predict gene co-functionality. These methods offer a powerful approach for predicting gene function, identifying disease genes, and reducing the search space of potential combinatorial gene effects to tractable levels. Coessentiality depends on some underlying variation in gene essentiality across the dataset; genes which are always essential or never essential rarely appear in these networks (Kim et al., 2019). In most approaches, the source of this variation is neither known nor questioned; however, the question is of high interest because identifying the causal basis for emergent essentiality would provide direct association of biomarkers with potential cancer targets as well as insight into the critical biological processes in different cells.

Another feature of raw coessentiality networks is that, as with prior efforts to build integrative functional interaction networks (Lee et al., 2004), they represent an integration of all contexts present in the cell lines or data sets from which they are derived. Functional interactions may often differ across contexts (Bandyopadhyay et al., 2010; Greene et al., 2015); for example, the roles of *FURIN* and *CPD* in promoting the maturation of different cell surface receptors in different tissues. Combining these contexts dilutes the overall correlation between genes with strong but context-dependent functional interaction. The approach we describe here offers a path towards understanding how context-dependent interactions can be gleaned from these data.

## Methods

### Preprocessing publicly available CRISPR screens

Raw read counts of CRISPR screens used in this study were downloaded from DepMap 20Q4 (n=808 cell lines). For DepMap screens, we removed guide RNAs targeting multiple protein coding genes based on the guide map obtained from the DepMap database and the CCDS protein coding gene annotation to avoid genetic interaction effects. After that, we applied the BAGEL2 pipeline (Kim and Hart, 2021) to measure gene essentiality information for each cell line. Gene knockout fitness phenotype is reported as a log Bayes Factor (BF), with positive scores indicating likelihood of essentiality.

### Predictive features and response variables for the logistic regression model

Genetic lesions were classified as gain or loss of function based on mutation calls described in the Cancer Cell Line Encyclopedia. For each of the 808 cell lines, each gene (n=18,111) was classified as loss of function (LOF) if it carried a lesion whose “Variant_annotation” annotation was ‘damaging’. Genes were characterized as gain of function (GOF) if they carried a lesion where either isTCGAhotspot or isCOSMIChotspot = True and if “Variant_annotation” was “other non-conserving.” Other lesions were not classified. See notebook “pre-step01-clean_data” for details.

CCLE Gene expression data in logTPM was downloaded from the CCLE portal at the Broad (n=718 cell lines with matching CRISPR data from Avana 20q4). EMT state was determined by CDH1/VIM expression log ratio (logTPM_CDH1_ – logTPM_VIM_), which resulted in a bimodal distribution across all cell lines. Each cell line was assigned CDH_VIM_hi if the log ratio > 1, or CDH_VIM_lo for log ratio < −4. See notebook “step01-merge_features” for details.

Other cell line metadata, downloaded from the CCLE portal at the Broad (n=808 cell lines with matching CRISPR data from Avana 20q4), were transformed into binary features. Each unique entry for “lineage”, “lineage_subtype”, “lineage_sub_subtype”, “culture_type”, and “sex” was considered a unique feature vector. See notebook “step01-merge_features” for details.

All features were merged into on matrix (808 cell lines x 36,406 features). Features describing six or fewer cell lines (n=33,475) were excluded. Remaining features were then de-duplicated by calculating and all-by-all correlation matrix and, for every pair of features with corr >= 0.9, the smaller feature (the one describing fewer cell lines) was discarded. For features with equal size and perfect overlap (e.g. “plasma_cell” and “multiple_myeloma”), one was selected at random and discarded. The final de-duplicated binary feature set comprised 2,918 predictor variables for the regression model. See notebook “step02-remove_duplicates” for details.

Gene essentiality was binarized for use as a response variable. Genes with missing values in the Bayes Factor table (n=17) were removed from the dataset. Then, for each cell line, genes with BF>=10 were classified as essential and all others nonessential, for a set of 18,094 genes in 808 cell lines. See notebook “step02-remove_duplicates” for details.

### Logistic regression model

Each gene’s binary essentiality vector was used as the response variable in a logistic regression model, using the binary features described above. Genes whose essentiality is invariant across the cell lines are uninteresting; therefore we excluded genes essential in less than 1% or more than 80% of the samples. The remaining set included 2,987 genes.

A logistic regression model was implemented in Python using sklearn.linear_model.LogisticRegression, with an elastic net penalty (L1 ratio = 0.25). Processing time took 78 minutes on a latest-generation AMD PC processor with 32 threads. The coefficients for each feature/gene prediction were combined into a single matrix. The intercept of the models closely matched the background probability of gene essentiality (Supplementary Figure 1). For each of the 2,987 response genes, the maximum and minimum feature coefficients were determined and, based on histograms of the maximum and minimum values (Supplementary Figure 1), |coeff| > 1.2 was chosen as a threshold for strong feature-gene association. The feature-gene coefficients matrix was flattened to a list and sorted by coefficient, and this list was used for subsequent analyses. A total of 393 unique features were predictive of gene essentiality with |coeff| > 1.2. See notebook “step03-do_regression” and “step03a-audit_regression” for details.

### Measurement of emergent network rewiring in CRISPR dataset

#### Leave-one-out test

To test for dynamic functional network rewiring, we measured the differential Pearson correlation coefficient (dPCC) between the global network, calculating for all gene pairs using all 808 cell lines (PCC_all_), and the network derived from a subset of cell lines that exclude a given context (PCC_∼context_). The formula of dPCC for each gene pair is therefore:

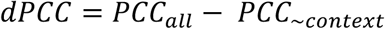

Positive dPCC represents a gain of interaction (GOI) because the higher PCC_all_ depends on the presence of cells with the feature of interest, and negative dPCC represents ‘loss of interaction’ because excluding cells associated with the feature of interest increases the PCC.

We evaluated the statistical significance of the dPCC distribution for each feature empirically. Briefly, we calculated the number of cell lines associated with the feature, ‘n’. Then we randomly selected ‘n’ cell lines from the set of all 808 cell lines, removed them, and calculated the all-by-all PCC matrix for all pairs of genes, PCC_rand_. For each gene pair, we measured PCC_all_ – PCC_rand_ for every gene. We repeated this process 1000x and compared dPCC to this distribution to generate an empirical P-value down to 0.001. We calculated this null distribution for every gene pair for each of the 393 contexts. See “calc_diff_coess_pval.py” for code and details.

## Acknowledgments

EK and TH were supported by NIGMS grant R35GM130119. EK was supported by a grant from the Prostate Cancer Foundation. TH is a CPRIT Scholar in Cancer Research (RR160032). This work was supported by the Andrew Sabin Family Foundation Fellowship and by NCI Cancer Center Support Grant P30CA16672.

## Author Contributions

EK developed the hypothesis. EK, TH, and VG performed bioinformatic analysis, with contributions from LN and CB. LN developed the diffnet web application under the supervision of CB. EK and TH drafted and all authors edited the manuscript.

---

**Figure S1.**
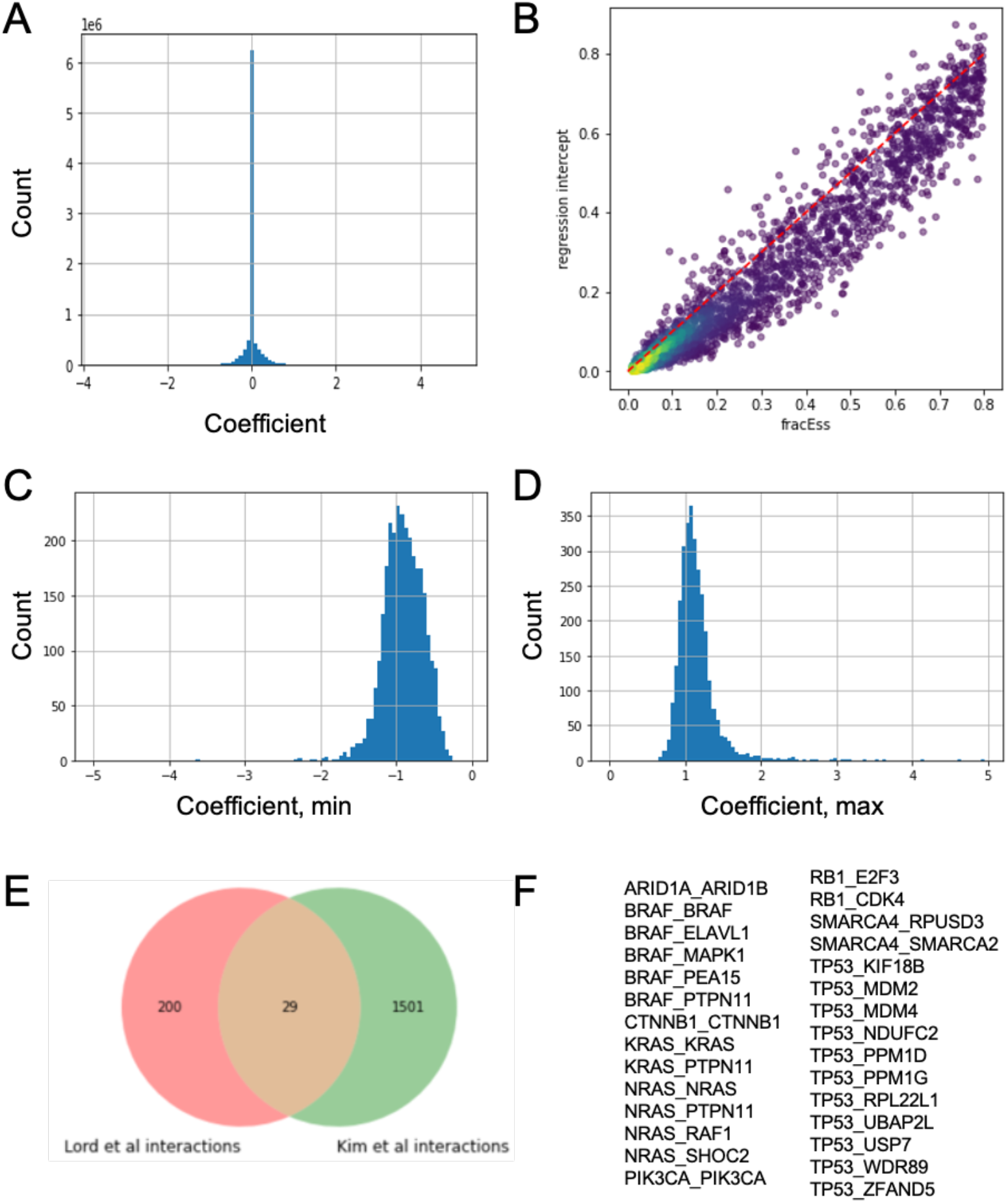
Regression model. (A) Histogram of all ∼8.7 million genomic feature-response gene coefficients. Elastic net penalty forces most coefficients to zero. (B) Regression intercept (y-axis, exp(Intercept) / exp(Intercept) + 1) approximates frequency of gene essentiality across cell lines. (C, D) Distribution of minimum (C) and maximum (D) value of regression coefficient for each of 2,918 predictor variables across 2,987 predictions, from which we chose |coeff| > 1.2 for further analysis. (E) Comparison of significant feature-gene interactions between Lord et al and this study. This study includes tissue/lineage as a predictor, resulting in more interactions. (F) Interactions (in mutated gene_response gene format) in the Lord/Kim intersection from (E).

## References

Aguirre, A.J., Meyers, R.M., Weir, B.A., Vazquez, F., Zhang, C.-Z., Ben-David, U., Cook, A., Ha, G., Harrington, W.F., Doshi, M.B., et al. (2016). Genomic Copy Number Dictates a Gene-Independent Cell Response to CRISPR/Cas9 Targeting. Cancer Discov 6, 914–929.

Alarcón, C., Cheatham, B., Lincoln, B., Kahn, C.R., Siddle, K., and Rhodes, C.J. (1994). A Kex2-related endopeptidase activity present in rat liver specifically processes the insulin proreceptor. Biochem J 301 (Pt 1), 257–265.

Amici, D.R., Jackson, J.M., Truica, M.I., Smith, R.S., Abdulkadir, S.A., and Mendillo, M.L. (2020). FIREWORKS: a bottom-up approach to integrative coessentiality network analysis. Life Sci Alliance 4, e202000882.

Aregger, M., Lawson, K.A., Billmann, M., Costanzo, M., Tong, A.H.Y., Chan, K., Rahman, M., Brown, K.R., Ross, C., Usaj, M., et al. (2020). Systematic mapping of genetic interactions for de novo fatty acid synthesis identifies C12orf49 as a regulator of lipid metabolism. Nature Metabolism 2, 499–513.

Asati, V., Mahapatra, D.K., and Bharti, S.K. (2016). PI3K/Akt/mTOR and Ras/Raf/MEK/ERK signaling pathways inhibitors as anticancer agents: Structural and pharmacological perspectives. Eur J Med Chem 109, 314–341.

Bandyopadhyay, S., Mehta, M., Kuo, D., Sung, M.-K., Chuang, R., Jaehnig, E.J., Bodenmiller, B., Licon, K., Copeland, W., Shales, M., et al. (2010). Rewiring of genetic networks in response to DNA damage. Science 330, 1385–1389.

Barretina, J., Caponigro, G., Stransky, N., Venkatesan, K., Margolin, A.A., Kim, S., Wilson, C.J., Lehár, J., Kryukov, G.V., Sonkin, D., et al. (2012). The Cancer Cell Line Encyclopedia enables predictive modelling of anticancer drug sensitivity. Nature 483, 603–607.

Behan, F.M., Iorio, F., Picco, G., Gonçalves, E., Beaver, C.M., Migliardi, G., Santos, R., Rao, Y., Sassi, F., Pinnelli, M., et al. (2019). Prioritization of cancer therapeutic targets using CRISPR–Cas9 screens. Nature 568, 511–516.

Boyle, E.A., Pritchard, J.K., and Greenleaf, W.J. (2018). High-resolution mapping of cancer cell networks using co-functional interactions. Mol. Syst. Biol. 14, e8594.

Cho, S.B., Kim, J., and Kim, J.H. (2009). Identifying set-wise differential co-expression in gene expression microarray data. BMC Bioinformatics 10, 109.

Ding, K.-F., Finlay, D., Yin, H., Hendricks, W.P.D., Sereduk, C., Kiefer, J., Sekulic, A., LoRusso, P.M., Vuori, K., Trent, J.M., et al. (2018). Network Rewiring in Cancer: Applications to Melanoma Cell Lines and the Cancer Genome Atlas Patients. Front Genet 9, 228.

Gkeka, P., Evangelidis, T., Pavlaki, M., Lazani, V., Christoforidis, S., Agianian, B., and Cournia, Z. (2014). Investigating the structure and dynamics of the PIK3CA wild-type and H1047R oncogenic mutant. PLoS Comput Biol 10, e1003895.

Gonzalez-Perez, A., Perez-Llamas, C., Deu-Pons, J., Tamborero, D., Schroeder, M.P., Jene-Sanz, A., Santos, A., and Lopez-Bigas, N. (2013). IntOGen-mutations identifies cancer drivers across tumor types. Nat Methods 10, 1081–1082.

Greene, C.S., Krishnan, A., Wong, A.K., Ricciotti, E., Zelaya, R.A., Himmelstein, D.S., Zhang, R., Hartmann, B.M., Zaslavsky, E., Sealfon, S.C., et al. (2015). Understanding multicellular function and disease with human tissue-specific networks. Nat Genet 47, 569–576.

Guan, Y., Gorenshteyn, D., Burmeister, M., Wong, A.K., Schimenti, J.C., Handel, M.A., Bult, C.J., Hibbs, M.A., and Troyanskaya, O.G. (2012). Tissue-specific functional networks for prioritizing phenotype and disease genes. PLoS Comput Biol 8, e1002694.

Han, K., Pierce, S.E., Li, A., Spees, K., Anderson, G.R., Seoane, J.A., Lo, Y.-H., Dubreuil, M., Olivas, M., Kamber, R.A., et al. (2020). CRISPR screens in cancer spheroids identify 3D growth-specific vulnerabilities. Nature 580, 136–141.

Hart, T., Chandrashekhar, M., Aregger, M., Steinhart, Z., Brown, K.R., MacLeod, G., Mis, M., Zimmermann, M., Fradet-Turcotte, A., Sun, S., et al. (2015). High-Resolution CRISPR Screens Reveal Fitness Genes and Genotype-Specific Cancer Liabilities. Cell 163, 1515–1526.

Helming, K.C., Wang, X., Wilson, B.G., Vazquez, F., Haswell, J.R., Manchester, H.E., Kim, Y., Kryukov, G.V., Ghandi, M., Aguirre, A.J., et al. (2014). ARID1B is a specific vulnerability in ARID1A-mutant cancers. Nature Medicine 20, 251–254.

Hsu, C.-L., Juan, H.-F., and Huang, H.-C. (2015). Functional Analysis and Characterization of Differential Coexpression Networks. Sci Rep 5, 13295.

Ideker, T., and Krogan, N.J. (2012). Differential network biology. Molecular Systems Biology 8, 565.

Janes, M.R., Zhang, J., Li, L.-S., Hansen, R., Peters, U., Guo, X., Chen, Y., Babbar, A., Firdaus, S.J., Darjania, L., et al. (2018). Targeting KRAS Mutant Cancers with a Covalent G12C-Specific Inhibitor. Cell 172, 578-589.e17.

Kim, E., and Hart, T. (2020). Improved analysis of CRISPR fitness screens and reduced off-target effects with the BAGEL2 gene essentiality classifier. BioRxiv 2020.05.30.125526.

Kim, E., and Hart, T. (2021). Improved analysis of CRISPR fitness screens and reduced off-target effects with the BAGEL2 gene essentiality classifier. Genome Med 13, 2.

Kim, E., Dede, M., Lenoir, W.F., Wang, G., Srinivasan, S., Colic, M., and Hart, T. (2019). A network of human functional gene interactions from knockout fitness screens in cancer cells. Life Sci. Alliance 2, e201800278.

Kim, J., Kim, I., Han, S.K., Bowie, J.U., and Kim, S. (2012). Network rewiring is an important mechanism of gene essentiality change. Sci Rep 2, 900.

Kim, W.K., Krumpelman, C., and Marcotte, E.M. (2008). Inferring mouse gene functions from genomic-scale data using a combined functional network/classification strategy. Genome Biology 9, S5.

Komada, M., Hatsuzawa, K., Shibamoto, S., Ito, F., Nakayama, K., and Kitamura, N. (1993). Proteolytic processing of the hepatocyte growth factor/scatter factor receptor by furin. FEBS Lett 328, 25–29.

Lavoie, H., and Therrien, M. (2015). Regulation of RAF protein kinases in ERK signalling. Nat Rev Mol Cell Biol 16, 281–298.

Lee, I., Date, S.V., Adai, A.T., and Marcotte, E.M. (2004). A probabilistic functional network of yeast genes. Science 306, 1555–1558.

Lord, C.J., Quinn, N., and Ryan, C.J. (2020). Integrative analysis of large-scale loss-of-function screens identifies robust cancer-associated genetic interactions. ELife 9, e58925.

Lui, T.W.H., Tsui, N.B.Y., Chan, L.W.C., Wong, C.S.C., Siu, P.M.F., and Yung, B.Y.M. (2015). DECODE: an integrated differential co-expression and differential expression analysis of gene expression data. BMC Bioinformatics 16, 182.

Martínez-Jiménez, F., Muiños, F., Sentís, I., Deu-Pons, J., Reyes-Salazar, I., Arnedo-Pac, C., Mularoni, L., Pich, O., Bonet, J., Kranas, H., et al. (2020). A compendium of mutational cancer driver genes. Nat Rev Cancer 20, 555–572.

Meyers, R.M., Bryan, J.G., McFarland, J.M., Weir, B.A., Sizemore, A.E., Xu, H., Dharia, N.V., Montgomery, P.G., Cowley, G.S., Pantel, S., et al. (2017). Computational correction of copy number effect improves specificity of CRISPR-Cas9 essentiality screens in cancer cells. Nat. Genet. 49, 1779–1784.

Noordermeer, S.M., Adam, S., Setiaputra, D., Barazas, M., Pettitt, S.J., Ling, A.K., Olivieri, M., Álvarez-Quilón, A., Moatti, N., Zimmermann, M., et al. (2018). The shieldin complex mediates 53BP1-dependent DNA repair. Nature 560, 117–121.

O’Leary, K. (2021). Tracing the origins of KRAS oncogene addiction. Nat Rev Cancer 21, 69.

Pan, J., Meyers, R.M., Michel, B.C., Mashtalir, N., Sizemore, A.E., Wells, J.N., Cassel, S.H., Vazquez, F., Weir, B.A., Hahn, W.C., et al. Interrogation of Mammalian Protein Complex Structure, Function, and Membership Using Genome-Scale Fitness Screens. Cell Systems.

Rancati, G., Moffat, J., Typas, A., and Pavelka, N. (2018). Emerging and evolving concepts in gene essentiality. Nat Rev Genet 19, 34–49.

Rauscher, B., Heigwer, F., Henkel, L., Hielscher, T., Voloshanenko, O., and Boutros, M. (2018). Toward an integrated map of genetic interactions in cancer cells. Mol. Syst. Biol. 14, e7656.

Santarpia, L., Lippman, S.M., and El-Naggar, A.K. (2012). Targeting the MAPK-RAS-RAF signaling pathway in cancer therapy. Expert Opin Ther Targets 16, 103–119.

Shalem, O., Sanjana, N.E., Hartenian, E., Shi, X., Scott, D.A., Mikkelson, T., Heckl, D., Ebert, B.L., Root, D.E., Doench, J.G., et al. (2014). Genome-scale CRISPR-Cas9 knockout screening in human cells. Science 343, 84–87.

Sharma, S., Dincer, C., Weidemüller, P., Wright, G.J., and Petsalaki, E. (2020). CEN-tools: an integrative platform to identify the contexts of essential genes. Mol Syst Biol 16.

Singh-Blom, U.M., Natarajan, N., Tewari, A., Woods, J.O., Dhillon, I.S., and Marcotte, E.M. (2013). Prediction and Validation of Gene-Disease Associations Using Methods Inspired by Social Network Analyses. PLOS ONE 8, e58977.

Terrell, E.M., and Morrison, D.K. (2019). Ras-Mediated Activation of the Raf Family Kinases. Cold Spring Harb Perspect Med 9, a033746.

Terrell, E.M., Durrant, D.E., Ritt, D.A., Sealover, N.E., Sheffels, E., Spencer-Smith, R., Esposito, D., Zhou, Y., Hancock, J.F., Kortum, R.L., et al. (2019). Distinct Binding Preferences between Ras and Raf Family Members and the Impact on Oncogenic Ras Signaling. Mol Cell 76, 872-884.e5.

Tsherniak, A., Vazquez, F., Montgomery, P.G., Weir, B.A., Kryukov, G., Cowley, G.S., Gill, S., Harrington, W.F., Pantel, S., Krill-Burger, J.M., et al. (2017). Defining a Cancer Dependency Map. Cell 170, 564-576.e16.

Wainberg, M., Kamber, R.A., Balsubramani, A., Meyers, R.M., Sinnott-Armstrong, N., Hornburg, D., Jiang, L., Chan, J., Jian, R., Gu, M., et al. (2021). A genome-wide atlas of co-essential modules assigns function to uncharacterized genes. Nat Genet 53, 638–649.

Wang, T., Birsoy, K., Hughes, N.W., Krupczak, K.M., Post, Y., Wei, J.J., Lander, E.S., and Sabatini, D.M. (2015). Identification and characterization of essential genes in the human genome. Science 350, 1096–1101.

Wang, T., Yu, H., Hughes, N.W., Liu, B., Kendirli, A., Klein, K., Chen, W.W., Lander, E.S., and Sabatini, D.M. (2017). Gene Essentiality Profiling Reveals Gene Networks and Synthetic Lethal Interactions with Oncogenic Ras. Cell 168, 890-903.e15.

Waters, A.M., and Der, C.J. (2018). KRAS: The Critical Driver and Therapeutic Target for Pancreatic Cancer. Cold Spring Harb Perspect Med 8, a031435.

Yu, C., Mannan, A.M., Yvone, G.M., Ross, K.N., Zhang, Y.-L., Marton, M.A., Taylor, B.R., Crenshaw, A., Gould, J.Z., Tamayo, P., et al. (2016). High-throughput identification of genotype-specific cancer vulnerabilities in mixtures of barcoded tumor cell lines. Nat Biotechnol 34, 419–423.

Zha, J., and Lackner, M.R. (2010). Targeting the insulin-like growth factor receptor-1R pathway for cancer therapy. Clin Cancer Res 16, 2512–2517.

Zimmermann, M., Murina, O., Reijns, M.A.M., Agathanggelou, A., Challis, R., Tarnauskaitė, Ž., Muir, M., Fluteau, A., Aregger, M., McEwan, A., et al. (2018). CRISPR screens identify genomic ribonucleotides as a source of PARP-trapping lesions. Nature 559, 285–289.

